# DAG-HEART: Directed Acyclic Graph-Guided Health Equity-Aware Representation Transfer Learning Framework for Breast Cancer

**DOI:** 10.64898/2026.08.31.748384

**Authors:** Minjeong Baek, Jieqiong Wang, Shibiao Wan

**Affiliations:** Department of Genetics, Cell Biology and Anatomy, University of Nebraska Medical Center, Omaha, NE, United States; Department of Neurological Sciences, University of Nebraska Medical Center, Omaha, NE, United States; Fred and Pamela Buffett Cancer Center, University of Nebraska Medical Center, Omaha, NE, United States

**Keywords:** Breast cancer, Directed acyclic graph, Kernel learning, Transfer learning, Data augmentation, Clinical outcome prediction, Cancer disparities

## Abstract

Breast cancer outcome prediction remains challenging for underrepresented populations because genomic datasets are demographically imbalanced and conventional multi-omics integration largely relies on undirected molecular similarity. We developed DAG-HEART, a directed acyclic graph-guided multi-omics transfer-learning framework that extends our previous transfer learning strategy with data augmentation. Using TCGA-BRCA mRNA, miRNA, and DNA-methylation data, DAG-HEART was evaluated for progression-free interval prediction in a data-minority group. DAG-guided nonlinear integration consistently improved predictive performance relative to direction-agnostic and correlation-based representations, while biologically motivated directional constraints generally outperformed reversed or unconstrained structures. Recurrently selected features converged on extracellular-matrix and regulatory pathways and supported clinically meaningful risk stratification. DAG-HEART provides an interpretable strategy for combining directed multi-omics structure with transfer learning under data imbalance across racial groups.

## Introduction

Breast cancer (BC) remains a major cause of cancer mortality among women in the United States, and substantial racial disparities persist despite advances in screening and treatment (Giaquinto et al., 2024; Siegel et al., 2025). Black women experience higher breast cancer mortality than White women and are disproportionately affected by aggressive disease phenotypes, although race itself should not be interpreted as an intrinsic biological determinant (Giaquinto et al., 2024; Newman & Kaljee, 2017). Race is a social construct shaped by historical, environmental, socioeconomic, and structural factors, whereas genetic ancestry reflects population history; conflating these concepts can lead to inappropriate biological interpretations of health disparities (Yudell et al., 2016; Borrell et al., 2021). At the same time, genomic resources remain demographically imbalanced, with European-ancestry populations overrepresented in cancer and human genetic studies, including large-scale sequencing initiatives such as The Cancer Genome Atlas (TCGA) (Spratt et al., 2016; Sirugo et al., 2019). Such imbalance creates a computational challenge because predictive models developed predominantly from majority populations may not generalize equally well to underrepresented populations, whereas limited target-population data constrain reliable model development (Gao & Cui, 2020; Sirugo et al., 2019). Therefore, improving prognostic reliability for underrepresented populations requires computational strategies that leverage information from larger cohorts without treating racial categories as biological mechanisms.

Transfer learning (TL) provides a principled strategy for this problem by transferring information learned from a data-majority source domain to a related but data-minority target domain and has been proposed specifically for mitigating performance disparities arising from biomedical data inequality (Pan & Yang, 2010; Gao & Cui, 2020). Data augmentation (DA) can complement this strategy by increasing the effective representation of limited target samples, although synthetic oversampling requires careful validation because generated samples may preserve biases or distort the original data distribution (Chawla et al., 2002; Shorten & Khoshgoftaar, 2019). We previously developed MOTLAB, which combined multi-omics integration with TL and DA for BC outcome prediction, building on established TL and minority-sample DA (Baek et al., 2025). However, MOTLAB represents patient similarity using Pearson correlation, a symmetric measure of association that cannot encode direction among multi-omics data (Rodgers & Nicewander, 1988; Schober et al., 2018). This limitation is relevant to multi-omics modeling because mRNA expression, miRNA abundance, and DNA methylation capture complementary molecular information, whereas widely used similarity- and latent-factor-based integration approaches primarily represent shared similarity or variation rather than directed dependencies (Cancer Genome Atlas Network, 2012; Hasin et al., 2017; Wang et al., 2014; Argelaguet et al., 2018). Directed acyclic graphs (DAGs) provide an alternative representation of directed dependencies, and continuous optimization methods have enabled efficient DAG structure learning (Zheng et al., 2018; Yu et al., 2019). Nevertheless, DAGs inferred from observational molecular data should not automatically be interpreted as causal networks because causal identification requires additional assumptions or interventional evidence (Pearl, 2009; Peters et al., 2017). These limitations motivate the integration of biologically informed direction constraints with multi-omics representation learning and TL for data-minority groups.

Here, we present DAG-HEART, a DAG-guided Health Equity-Aware Representation TL framework that extends the TL and DA strategy established in MOTLAB by introducing direction-constrained multi-omics representation learning. DAG-HEART reduces mRNA, miRNA, and DNA-methylation profiles into latent components, estimates directed dependencies among these components under biologically motivated block-level constraints, and uses the resulting structure to guide nonlinear patient-similarity kernels. Importantly, the learned graph is interpreted as a structured representation of directed statistical dependencies rather than evidence of gene-level causation. Building on MOTLAB, DAG-HEART transfers information from the data-majority group to the smaller target population while incorporating DA of data-minority group during model development. We evaluated DAG-HEART using TCGA-BRCA multi-omics data and standardized progression-free interval definitions derived from established TCGA resources (Cancer Genome Atlas Network, 2012; Liu et al., 2018). Prognostic performance was assessed at 2-, 3-, 4-, and 5-year PFI time settings using repeated cross-validation and compared with Pearson-correlation-based integration and alternative DAG direction constraints. We further examined recurrently selected molecular features and pathway-level convergence to assess whether predictive signals were accompanied by interpretable molecular patterns. Thus, DAG-HEART is the integration of biologically constrained directional multi-omics structure with nonlinear patient-similarity learning and the TL and DA strategy of MOTLAB, providing a framework for clinical outcome prediction to reduce BC disparities.

## Results

### DAG-HEART integrates direction-constrained multi-omics representations with transfer learning for reducing breast cancer disparities

To determine whether directed multi-omics structure could improve clinical outcome prediction under demographic data imbalance, we developed DAG-HEART as an extension of our previous TL and DA strategy. Female patients from TCGA-BRCA were partitioned into a European American (EA) group and an African American (AA) group, and progression was predicted independently at 2-, 3-, 4-, and 5-year progression-free interval (PFI) settings (**Fig. 1A**). TCGA-BRCA provides complementary molecular measurements across transcriptional, post-transcriptional, and epigenetic layers, and PFI has been systematically curated as a clinical outcome for TCGA survival analyses (Cancer Genome Atlas Network, 2012; Liu et al., 2018).

**Figure 1.**
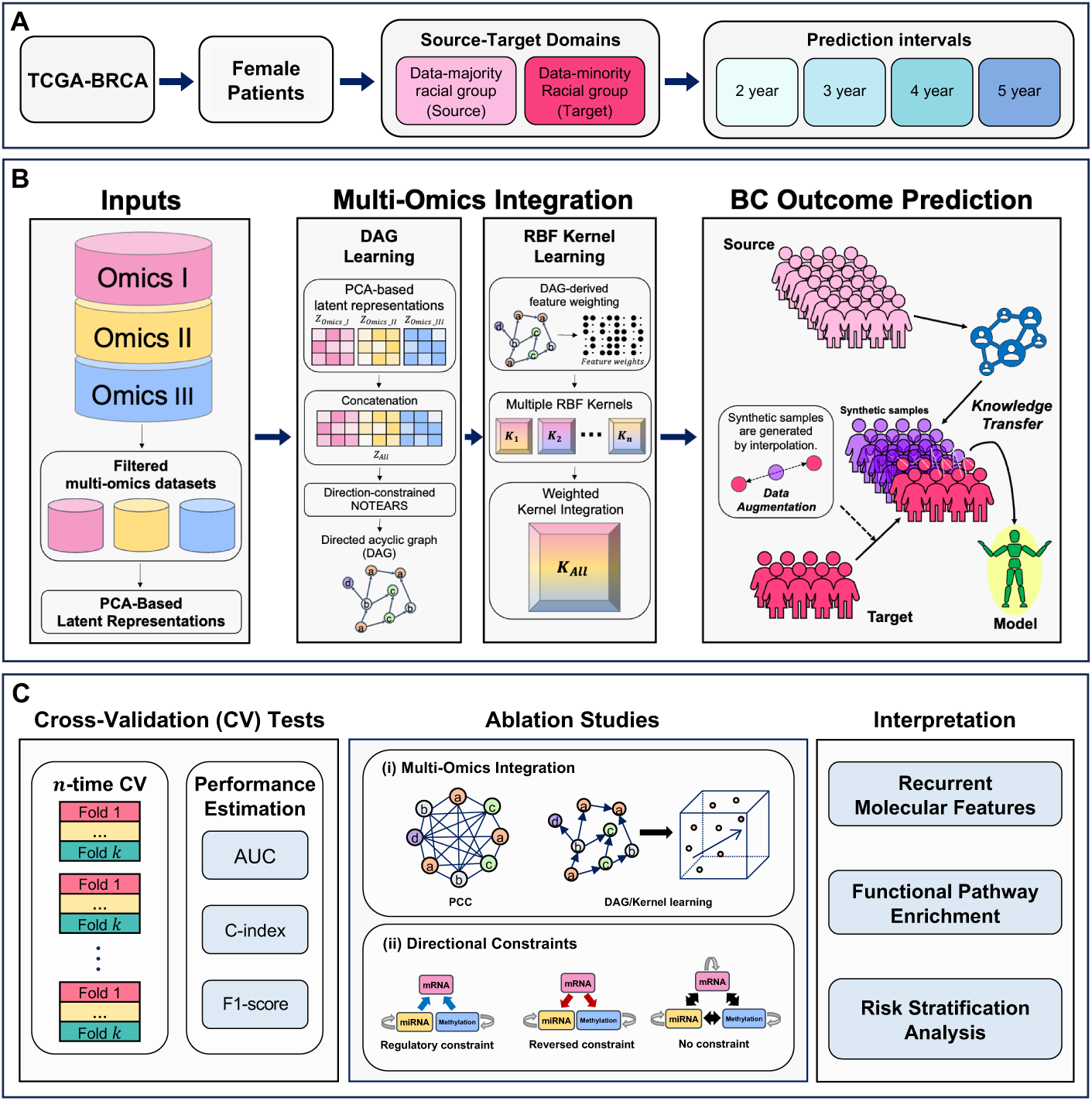
Overview of the DAG-HEART framework and evaluation strategy. (A) TCGA-BRCA study design showing source and target populations and 2-, 3-, 4-, and 5-year progression-free interval (PFI) prediction tasks. (B) Multi-omics integration of mRNA, miRNA, and DNA methylation using PCA-based latent representations, direction-constrained DAG learning, DAG-weighted RBF kernels, multiple-kernel integration, and transfer learning with target-domain augmentation. (C) Evaluation using 10 repeated three-fold cross-validations, architectural and directional ablation analyses, recurrent molecular-feature analysis, pathway enrichment, and risk stratification.

DAG-HEART incorporated mRNA expression, miRNA expression, and DNA methylation through two sequential representation-learning stages (**Fig. 1B**). After feature filtering and principal-component reduction, latent representations from the three omics layers were jointly modeled using direction-constrained DAG learning. The resulting directed structure was then incorporated into radial-basis-function (RBF) patient-similarity kernels, followed by multiple-kernel integration and nonlinear dimensionality reduction. Kernel methods provide a principled mechanism for representing nonlinear similarity, whereas multiple-kernel learning enables complementary similarity structures to be combined across heterogeneous information sources (Schölkopf et al., 1998; Gönen & Alpaydın, 2011). The learned representation was subsequently coupled to knowledge transfer from the source domain and target-domain DA before prediction in the target domain.

The evaluation design was constructed to distinguish predictive improvement from methodological artifacts. Models were assessed using ten repeats of three-fold cross-validation with AUROC, weighted F1 score, C-index, and PR-AUC as complementary performance measures (**Fig. 1C**). In parallel, ablation analyses separately tested the contributions of multi-omics integration and directional constraints, while downstream analyses examined recurrent molecular features, pathway enrichment, and risk stratification. This design allowed us to test not only whether DAG-HEART improved prediction, but also which component of the framework generated the improvement and whether the resulting representation retained molecular and clinical interpretability.

### Directed acyclic graph-guided nonlinear integration drives the predictive advantage of DAG-HEART

To address the contributions of DAG and nonlinear representation, we performed ablation study under four PFI settings (**Fig. 2A**). The baseline linear model used Pearson correlation coefficient (PCC)-based integration without DAG information, whereas its DAG-guided counterpart incorporated DAG-derived information into the same linear representation. In parallel, nonlinear models combined RBF kernels, multiple-kernel learning, and kernel PCA either without or with DAG guidance. This factorial comparison separated the effect of directionality from that of nonlinear patient-similarity learning.

**Figure 2.**
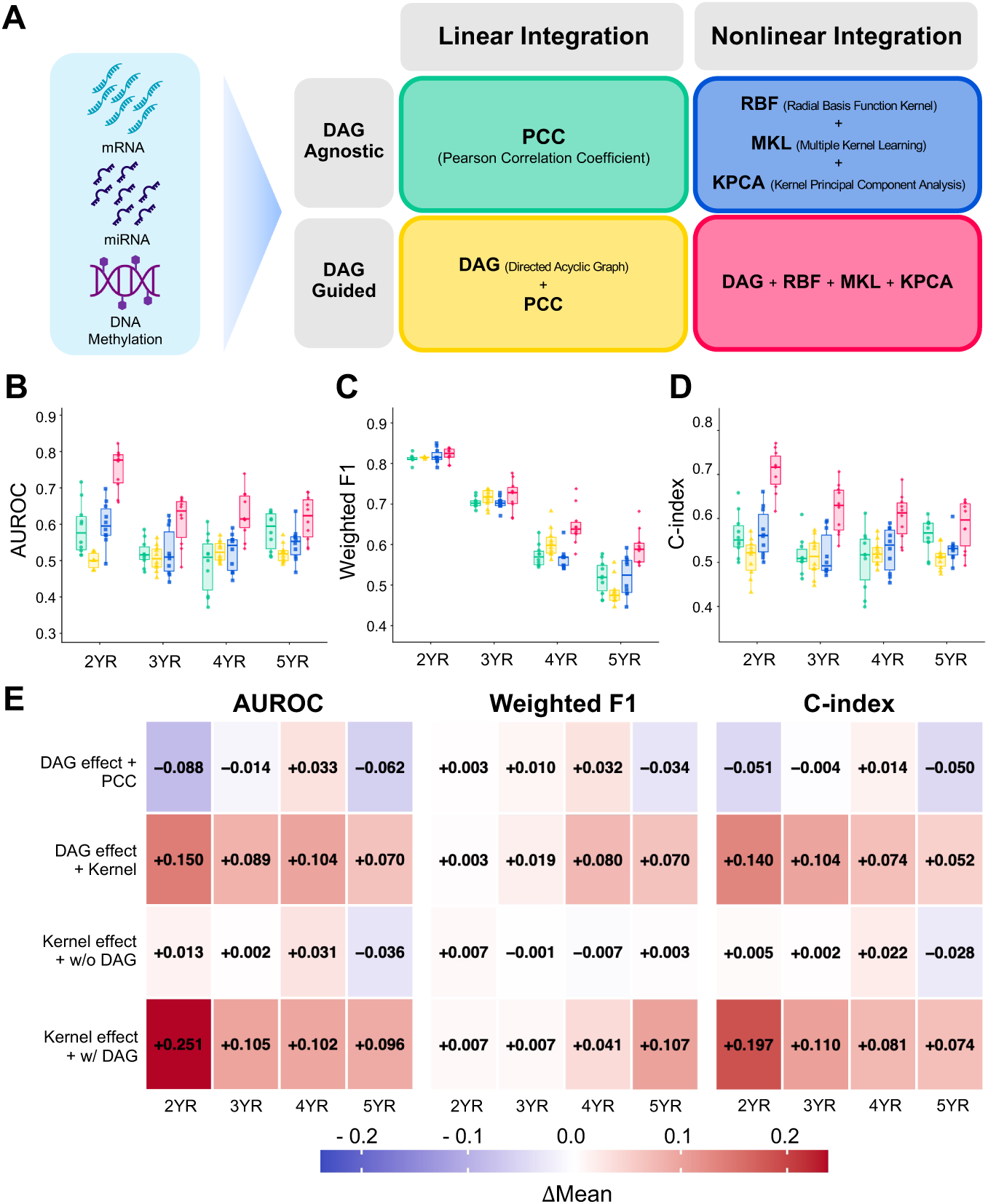
Architectural ablation of directed and nonlinear multi-omics integration. (A) Four integration architectures combining DAG-agnostic or DAG-guided representations with linear PCC or nonlinear RBF–MKL–KPCA integration. (B–D) Cross-validation distributions of AUROC, weighted F1, and C-index, respectively, across 2-, 3-, 4-, and 5-year PFI tasks. (E) Mean paired performance changes attributable to DAG guidance or nonlinear kernel integration. Positive values indicate improvement after addition of the indicated component.

Across the four PFI settings, the complete DAG-guided nonlinear model showed the most favorable overall distributions for AUROC, weighted F1, and C-index (**Fig. 2B–D**). Importantly, the component-wise analysis demonstrated that neither DAG information nor nonlinear integration alone accounted for this pattern (**Fig. 2E**). Adding DAG guidance to the PCC-based linear representation produced mean AUROC changes of −0.088, −0.014, +0.033, and −0.062 at 2, 3, 4, and 5 years, respectively, with similarly mixed changes in C-index. Thus, simply imposing directed information on a linear correlation representation did not consistently improve prediction. In contrast, the effect of DAG guidance became consistently positive when applied within the nonlinear kernel representation. Relative to the corresponding DAG-agnostic kernel model, DAG guidance increased mean AUROC by +0.150, +0.089, +0.104, and +0.070 across the four PFI settings and increased mean C-index by +0.140, +0.104, +0.074, and +0.052, respectively (**Fig. 2E**). Weighted F1 changes were also positive across all four PFI settings. The reciprocal comparison produced an even clearer pattern. Adding nonlinear kernel integration in the presence of DAG guidance increased mean AUROC by +0.251, +0.105, +0.102, and +0.096, whereas adding the same kernel component without DAG guidance resulted in only small or inconsistent changes. The corresponding C-index gains with DAG guidance were +0.197, +0.110, +0.081, and +0.074. These ablations identify an important property of the framework. The predictive contribution of directed multi-omics structure depended on the representation in which that structure was expressed. Directionality was not independently beneficial when superimposed on PCC-based linear integration; rather, its advantage emerged when directed relationships were translated into nonlinear patient similarities.

We further asked whether this improvement resulted merely from imposing any directed graph or specifically from the biologically motivated orientation of the omics layers. The regulatory constraint, reverse constraint, and unconstrained DAG formulations were therefore evaluated separately (**Fig. 3A**). The regulatory orientation generally produced more favorable AUROC, weighted F1, and C-index than the reversed orientation (**Fig. 3B–D**) and similarly tended to outperform the unconstrained formulation (**Fig. 3E–G**), although the magnitude of the difference varied by all PFI settings and metric. These results argue against graph direction serving only as an additional source of model flexibility. Instead, the orientation of cross-omics dependencies itself contributed to predictive performance, supporting the use of biologically informed directional structure as a representation prior rather than an arbitrary graph constraint.

**Figure 3.**
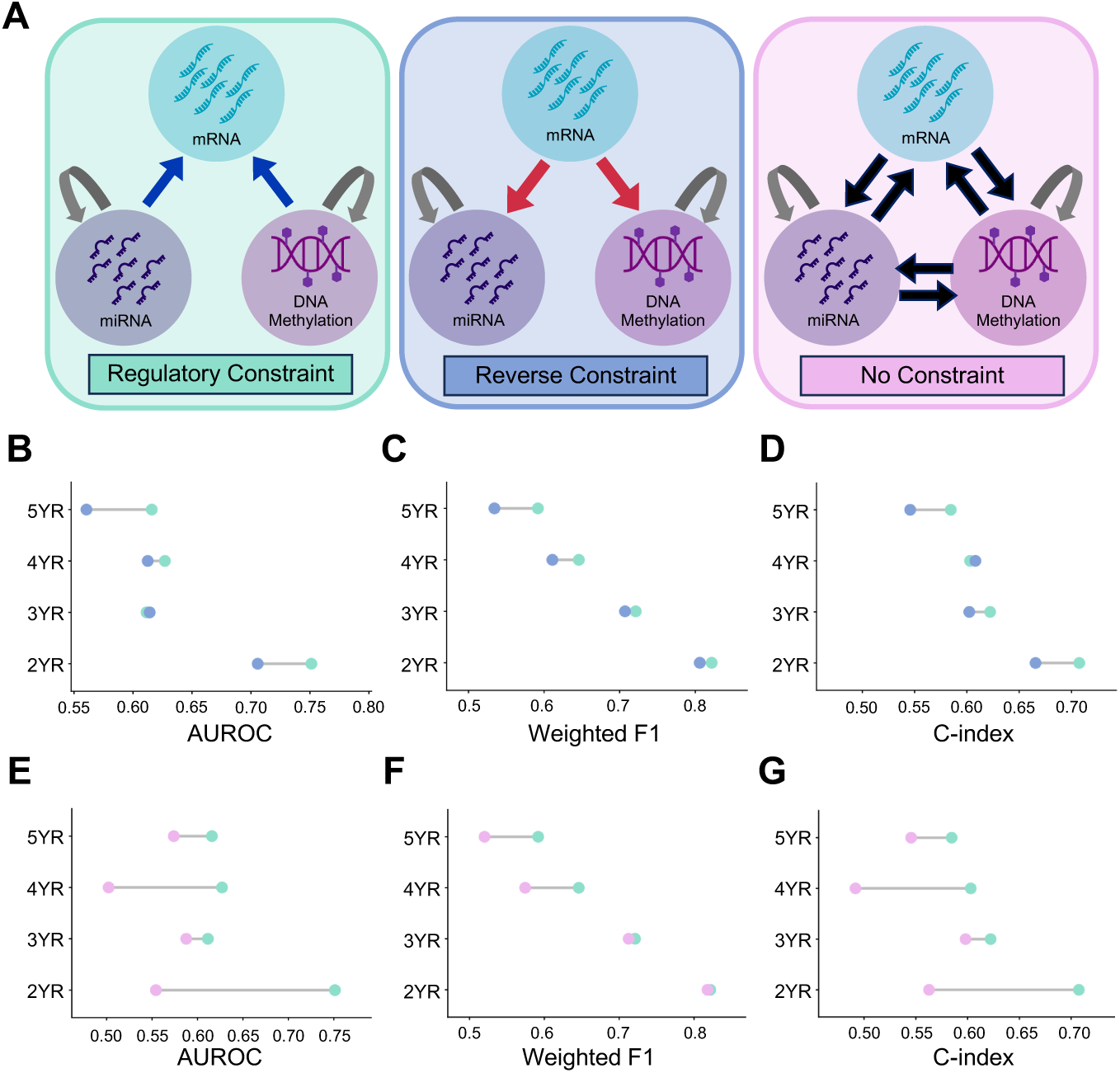
Effect of biological direction constraints on DAG-HEART performance. (A) DAG structures evaluated using the proposed regulatory constraint, reversed cross-omics directions, or no block-level directional constraint. (B–D) Performance comparison between the regulatory and reversed constraints for AUROC, weighted F1, and C-index across 2–5-year PFI tasks. (E–G) Corresponding comparison between the regulatory and unconstrained models. Connected points represent matched prediction horizons. The analyses assess whether predictive performance depends on the biologically motivated orientation of cross-omics dependencies rather than DAG structure alone.

### DAG-HEART captured stable molecular features reveal cross-omics convergence on extracellular-matrix and regulatory programs

Having established the predictive contribution of DAG-guided integration, we next examined whether the models repeatedly selected reproducible molecular features across PFI settings. Feature recurrence was evaluated separately for mRNA, miRNA, and gene-level DNA methylation, revealing subsets with both high mean importance and frequent selection across repeated PFI prediction tasks (**Fig. 4A–C**). Several mRNA features, including SPIN4, members of the SNORD114 family, MIR599, F13B, CSN2, and CDH19, showed high recurrence, while recurrent miRNA signals included miR-192, miR-204, miR-27a, miR-23a, miR-215, and miR-194 family members (**Fig. 4D**). Recurrent methylation-associated features included P4HA3, FAP, DACT1, COL6A3, COL5A2, COL5A1, CHSY3, CDH11, and TVP23C. Importantly, the stability profiles were not restricted to a single PFI prediction. Subsets of recurrent features were shared across multiple PFI settings, particularly within the mRNA and DNA-methylation layers (**Fig. 4E**).

**Figure 4.**
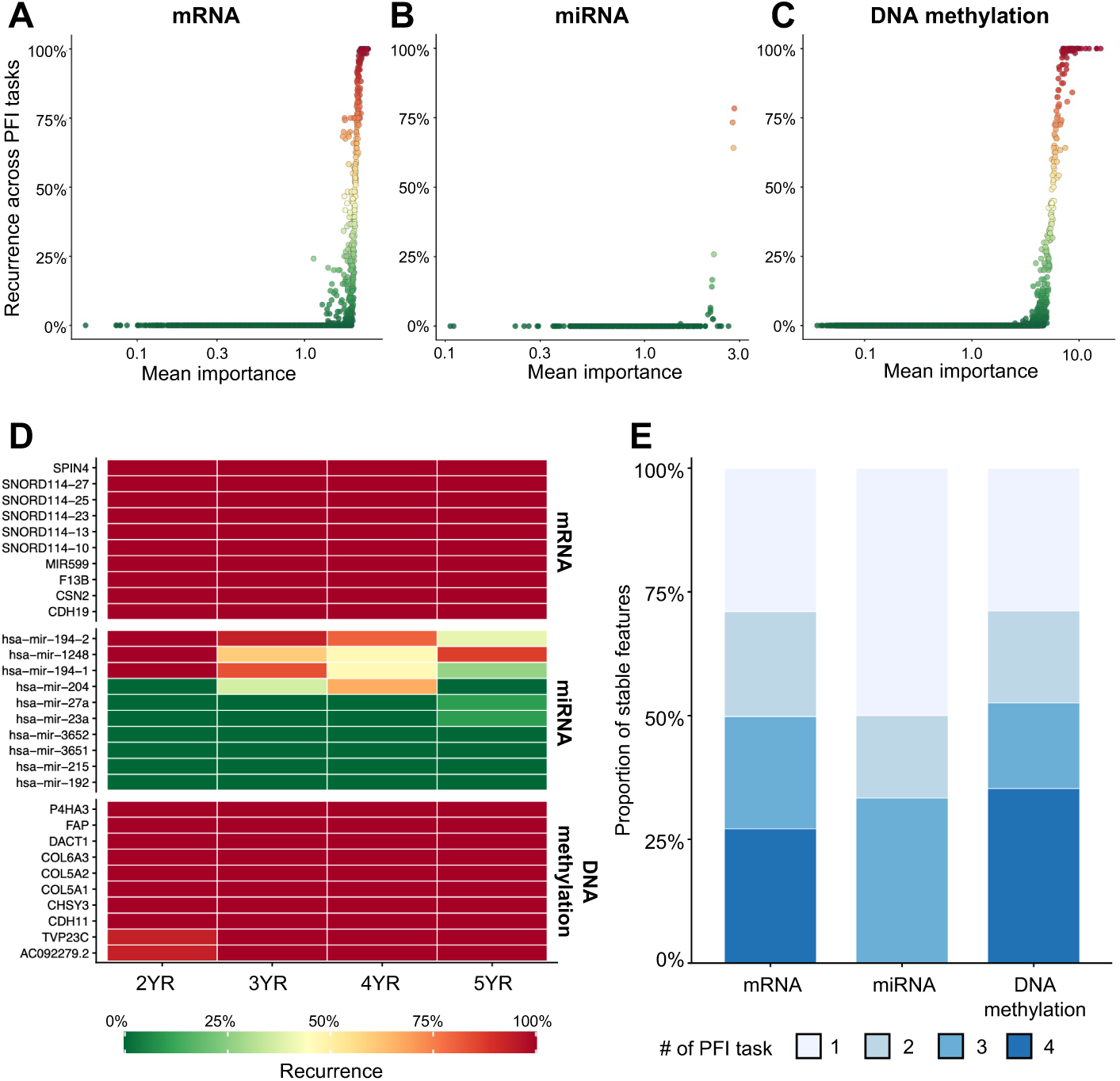
Stability of recurrent molecular features across PFI prediction tasks. (A–C) Mean structural importance versus recurrence across repeated models for mRNA, miRNA, and DNA-methylation features, respectively. Recurrence represents the frequency with which a feature was repeatedly identified among highly ranked features. (D) Recurrence of representative stable features across the 2-, 3-, 4-, and 5-year PFI tasks. (E) Proportion of stable features recurring across one to four prediction horizons within each omics layer. Structural importance was derived from the DAG-guided representation and does not indicate causal effects.

Pathway analysis of the stable features revealed distinct but complementary biological patterns across omics layers (**Fig. 5**). The mRNA-derived signal was dominated by extracellular-matrix organization, collagen formation, ECM proteoglycans, collagen biosynthesis and modification, matrix degradation, integrin interactions, and collagen crosslinking (**Fig. 5A**). This pattern is biologically coherent with extensive evidence that remodeling of the extracellular matrix and collagen architecture affects tumor-cell behavior, invasion, and cancer progression (Cox & Erler, 2011; Pickup et al., 2014).

**Figure 5.**
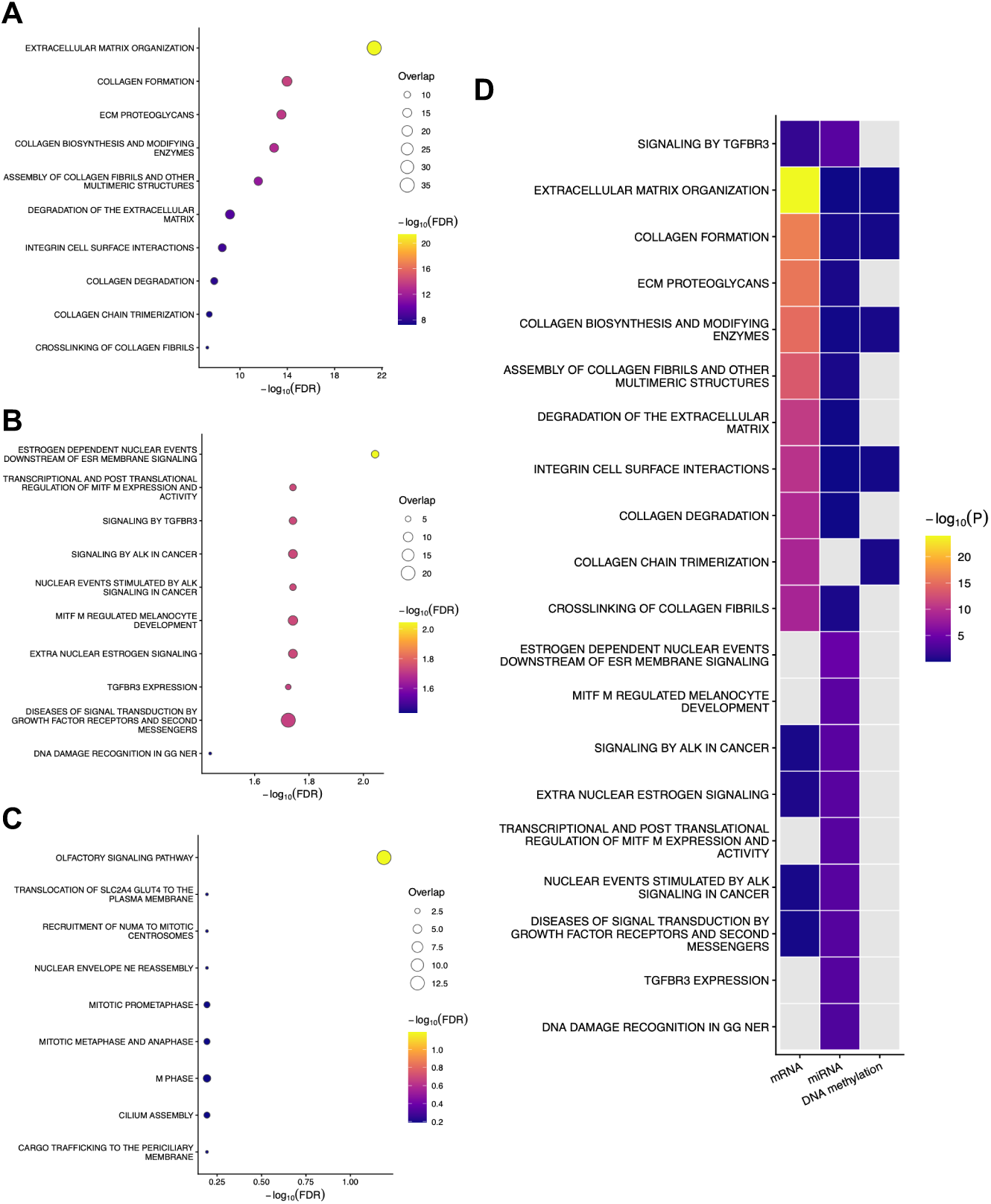
Functional pathway enrichment of stable DAG-HEART molecular features. (A–C) Reactome over-representation analysis for stable mRNA features, experimentally validated targets of stable miRNAs, and stable DNA-methylation-associated genes, respectively. Point position and color represent enrichment significance, and point size represents the number of overlapping genes. (D) Cross-omics comparison of pathways identified across the three molecular layers; color intensity represents nominal enrichment strength. Pathway enrichment was used for functional interpretation and does not establish mechanistic or causal involvement.

miRNA-associated targets showed a different enrichment profile, including estrogen-dependent nuclear signaling, MITF-related transcriptional regulation, TGFBR3 expression processes, and ALK-associated signaling (**Fig. 5B**). DNA-methylation-associated features generated comparatively weaker pathway-level evidence, with signals involving cell-cycle, nuclear-envelope, trafficking, and related processes (**Fig. 5C**); these results were therefore interpreted more cautiously than the stronger mRNA enrichment. Cross-omics comparison further demonstrated that pathway support was heterogeneous rather than artificially uniform across all three modalities (**Fig. 5D**). Overall, these analyses indicate that the recurrent features captured by DAG-HEART were not simply a collection of individually high-importance variables, but converged most strongly on coherent extracellular-matrix and regulatory programs while retaining omics-specific signals.

### DAG-HEART risk scores stratify progression risk across PFI prediction settings and molecular subtypes

Finally, we evaluated whether the learned target-population predictions translated into clinically interpretable progression-risk stratification. At each of PFI settings, patients classified as high risk showed lower progression-free probabilities than those classified as low risk (**Fig. 6A**). Consistent with these survival curves, hazard-ratio point estimates comparing high- and low-risk groups were greater than one across all four models (**Fig. 6B**). The precision of these estimates varied across PFI settings, with the 3-year confidence interval extending across the null while the separation was clearer for several of the other prediction intervals. Thus, the prognostic signal was directionally consistent across independently PFI prediction settings rather than being restricted to a single setting.

**Figure 6.**
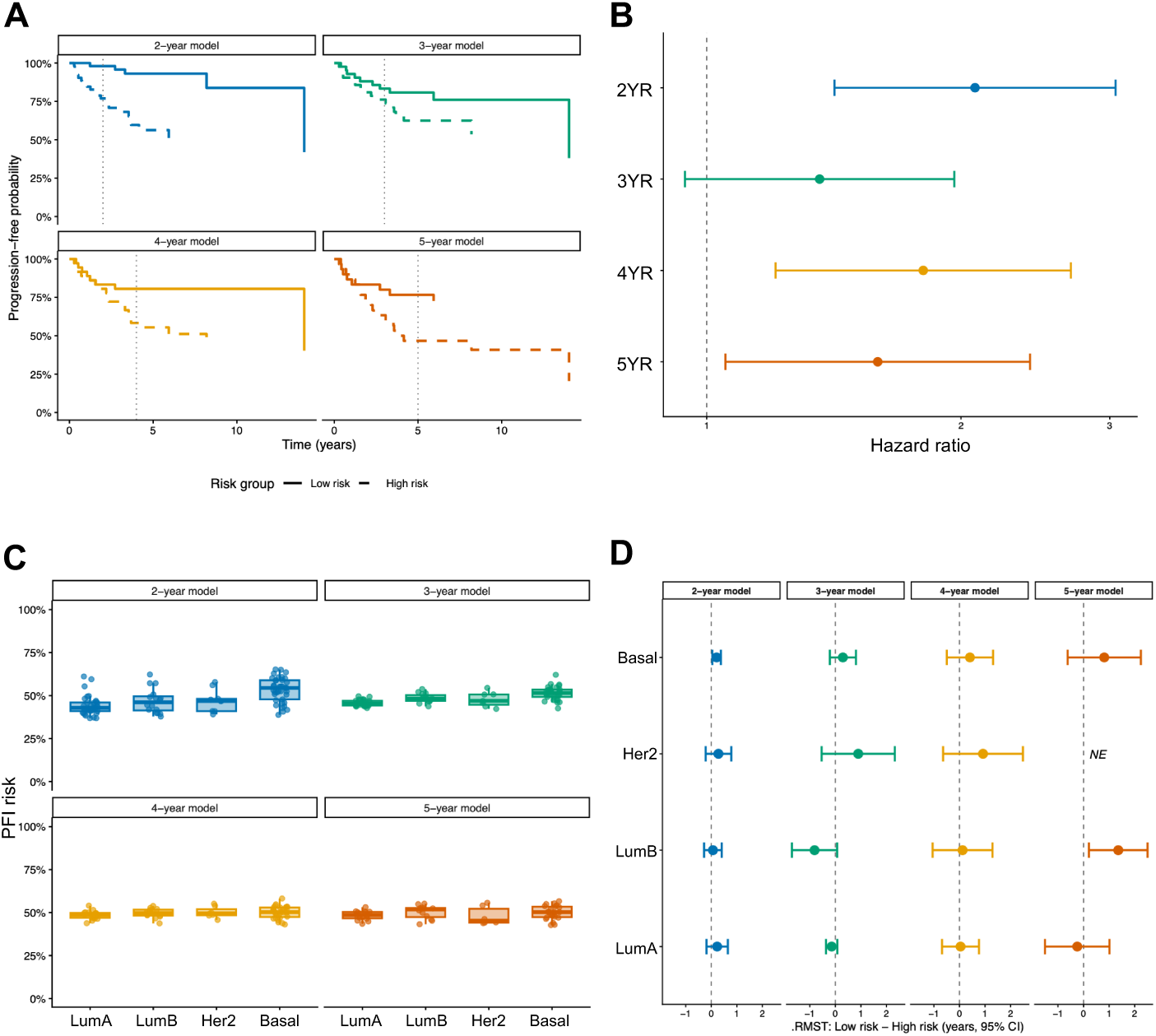
Clinical interpretation of DAG-HEART-derived progression risk. (A) Kaplan–Meier progression-free curves for model-defined low- and high-risk groups from the 2-, 3-, 4-, and 5-year PFI models. (B) Hazard ratios with 95% confidence intervals for continuous DAG-HEART progression risk. (C) Distribution of predicted PFI risk across PAM50 intrinsic subtypes for each prediction horizon. (D) Within-subtype differences in restricted mean survival time (RMST), calculated as low-risk minus high-risk survival time with 95% confidence intervals. Positive RMST differences favor the low-risk group; NE denotes not estimable.

We next examined the relationship between DAG-HEART risk and PAM50 intrinsic subtype. PAM50 captures major transcriptionally defined breast-cancer subtypes with established differences in tumor biology and prognosis (Perou et al., 2000; Parker et al., 2009). Predicted PFI risk varied among basal-like, HER2-enriched, Luminal B, and Luminal A tumors, but substantial within-subtype variability and overlap remained across all PFI predictions (**Fig. 6C**). This pattern indicates that DAG-HEART risk was not simply a surrogate for assigning tumors to established intrinsic subtypes. Consistently, restricted mean survival time analyses comparing model-defined low- and high-risk groups within PAM50 strata produced positive low-risk minus high-risk estimates across most estimable subtype–PFI setting combinations (**Fig. 6D**), although confidence intervals were wider in several subgroup analyses and the 5-year HER2-enriched estimate was not estimable. These subgroup results should therefore be interpreted as evidence of clinical relevance rather than definitive subtype-specific validation.

## Discussion

Breast cancer outcome prediction in demographically imbalanced genomic datasets presents two interconnected challenges. Molecular heterogeneity must be represented across multiple regulatory layers, while models must also remain informative for populations represented by substantially fewer training samples. Existing multi-omics approaches, including similarity network fusion and latent-factor models, have established the value of combining complementary molecular measurements, whereas TL provides a framework for leveraging data-majority groups to improve learning in data-minority groups (Wang et al., 2014; Argelaguet et al., 2018; Gao & Cui, 2020). Building on this landscape and on the TL and DA strategy previously implemented in MOTLAB, we developed DAG-HEART to incorporate biologically constrained directionality into multi-omics patient representation. Rather than treating mRNA, miRNA, and DNA methylation as interchangeable sources of undirected similarity, DAG-HEART learns directed dependencies among their latent representations and propagates this structure into nonlinear patient-similarity kernels before cross-population knowledge transfer. The principal contribution of this design is therefore not simply the addition of a DAG or another multi-omics integration step, but the coupling of direction-constrained molecular structure, nonlinear patient representation, and learning under target-population data scarcity within a single prognostic framework. Our ablation results clarify why this distinction matters. DAG guidance alone did not consistently improve a linear PCC-based representation, whereas incorporating the same directional information into nonlinear kernel learning produced favorable changes in AUROC, weighted F1, and C-index across PFI settings. Moreover, the biologically motivated regulatory constraint generally performed more favorably than reversed or unconstrained graph formulations. Together, these findings suggest that directional information becomes useful not merely by increasing model complexity, but when it shapes the geometry through which molecular similarity between patients is represented. This interpretation distinguishes DAG-HEART from conventional similarity-based integration and supports biologically informed directionality as a representation prior rather than as a claim of causal regulatory discovery. This distinction is essential because DAGs inferred from observational data encode statistical structure but do not establish causation without substantially stronger assumptions or interventional evidence (Pearl, 2009; Peters et al., 2017).

The downstream analyses further suggest that the predictive advantage of DAG-HEART was accompanied by molecular structure that was reproducible across related prognostic tasks rather than being confined to isolated high-importance features. Recurrently selected features were observed across mRNA, miRNA, and DNA-methylation layers, with the strongest pathway-level convergence emerging for extracellular-matrix organization, collagen formation and remodeling, ECM proteoglycans, and integrin-associated processes. These findings are biologically plausible because extracellular-matrix composition, collagen remodeling, and tumor–stroma interactions are established components of cancer invasion and progression, although enrichment in the present study should be interpreted as associative rather than mechanistic evidence (Cox & Erler, 2011; Pickup et al., 2014). The unequal strength of pathway enrichment across omics layers is also informative. In particular, the weaker DNA-methylation enrichment compared with the pronounced mRNA-associated ECM signal argues against interpreting the multi-omics analysis as requiring identical biological information from every modality; instead, complementary modalities may contribute differently to a shared predictive representation. At the clinical level, DAG-HEART-derived risk groups showed separation in progression-free outcomes across multiple PFI predictions, while predicted risk remained heterogeneous within established PAM50 subtypes. PAM50 subtypes capture major transcriptional distinctions among breast tumors and have established prognostic relevance (Perou et al., 2000; Parker et al., 2009). Thus, the persistence of within-subtype risk variation suggests that the learned multi-omics representation was not merely reproducing intrinsic subtype assignment. The generally positive restricted mean survival-time differences between low- and high-risk groups within subtype strata provide additional support for this interpretation, although several confidence intervals were wide and one subtype–PFI setting combination was not estimable. Accordingly, these findings should not yet be interpreted as demonstrating independent clinical utility beyond PAM50. Rather, they provide convergent evidence that the DAG-guided representation captures prognostically relevant information at multiple levels. This progression from model decomposition to molecular and clinical interpretation is important because performance improvements in high-dimensional multi-omics models are considerably more compelling when the source of the improvement can be interrogated and the resulting signals can be related to established tumor biology.

Several limitations define important directions for extending this work. First, DAG-HEART was developed and evaluated within TCGA-BRCA using repeated cross-validation; although repeated resampling provides a more robust assessment of internal performance than a single train–test split, it does not substitute for validation in a fully independent cohort. TCGA was designed primarily as a molecular characterization resource rather than a prospective prognostic study, and heterogeneity in follow-up and clinical annotation remains relevant even when standardized TCGA Clinical Data Resource endpoints are used (Cancer Genome Atlas Network, 2012; Liu et al., 2018). Independent validation should therefore be a priority, ideally using multi-institutional cohorts containing comparable mRNA, miRNA, DNA-methylation, outcome, and population annotations, followed ultimately by prospective evaluation. Second, the smaller target population necessarily limits the precision of subgroup analyses, including PAM50-stratified survival comparisons, and synthetic DA increases representation during model training but does not create independent biological observations. Larger enrollment of historically underrepresented populations remains the more fundamental solution to biomedical data inequality; TL should complement rather than replace efforts to improve representation in genomic studies (Gao & Cui, 2020; Sirugo et al., 2019). Third, the direction-constrained DAG was learned among dimension-reduced omics components under block-level regulatory priors. These edges should therefore be interpreted as directed statistical dependencies among latent representations, not as gene-specific regulatory interactions or causal effects. Furthermore, block-level constraints necessarily simplify context-dependent relationships among DNA methylation, miRNA regulation, and transcription, and future models could incorporate experimentally supported regulatory priors, additional molecular layers, or perturbational data to refine these structures. Fourth, grouping patients into source and target populations addresses a specific form of data imbalance but cannot disentangle genetic ancestry from environmental, socioeconomic, treatment-related, and structural determinants associated with racial disparities; race should not be interpreted as a biological mechanism (Yudell et al., 2016; Borrell et al., 2021). Finally, the recurrent-feature and pathway analyses remain hypothesis-generating and require replication in independent molecular datasets and experimental validation before individual genes or pathways can be considered mechanistically implicated in progression. Future work that combines larger and more diverse cohorts, external and prospective validation, richer clinical and treatment variables, refined biological constraints, calibration and clinical-utility assessment, and functional validation of recurrent molecular programs would provide a stronger test of generalizability and translational value. Within these boundaries, DAG-HEART establishes a foundation for studying whether biologically structured multi-omics representations can be combined with TL to improve clinical outcome prediction when clinically relevant populations are represented unequally in biomedical data.

## Materials and Methods

### Study cohort and multi-omics data

Multi-omics and clinical data from The Cancer Genome Atlas Breast Invasive Carcinoma cohort (TCGA-BRCA) were used in this study. Three molecular data types were analyzed, mRNA expression, miRNA expression, and DNA methylation. Samples were matched across the three omics datasets using the first 12 characters of the TCGA participant barcode, and only patients represented in all three molecular datasets were retained. Molecular features with measurements available in fewer than 95% of samples or with zero variance were removed, and remaining missing values were imputed using the feature-wise median. The three omics matrices were subsequently aligned to a common patient order.

Genetic-ancestry annotations were obtained from the TCGA genetic-ancestry resource, in which global ancestry was inferred using EIGENSTRAT-based population-structure analysis (Yuan et al., 2018). Patients assigned to European ancestry (EA) and African ancestry (AA) were included in the analysis, whereas other ancestry groups were excluded. EA patients constituted the data-rich source domain and AA patients constituted the underrepresented target domain. Genetic ancestry was used to define computational source and target domains and was not interpreted as self-identified race or as a causal biological determinant of breast cancer outcome. The genetic-ancestry resource used here was developed from genome-wide genotyping information across TCGA cancers and distinguishes ancestry from clinical race/ethnicity annotations (Yuan et al., 2018).

Progression-free interval (PFI) status and follow-up times were obtained from the harmonized TCGA Pan-Cancer Clinical Data Resource (TCGA-CDR; Liu et al., 2018). PFI prediction was evaluated separately at 2-, 3-, 4-, and 5-year PFI settings. For each PFI setting, patients censored at or before the corresponding time point were excluded because their fixed-time progression status could not be determined. Among the remaining patients, individuals who experienced a PFI event at or before the specified time were classified as having progression, whereas patients remaining progression-free beyond the time constituted the progression-free group. In the implemented classifier, progression-free status beyond the prediction time was encoded as the positive class. The TCGA-CDR provides harmonized survival outcomes specifically developed to support consistent outcome analyses across TCGA tumor types (Liu et al., 2018).

### Direction-constrained multi-omics representation learning

Within each training fold, the mRNA, miRNA, and DNA-methylation blocks were standardized separately and reduced by principal component analysis. Up to 16 principal components were retained from each omics block, with the number further limited by the available number of features and training samples. The resulting latent representations were concatenated, and a directed acyclic graph (DAG) was estimated over the multi-omics latent components using a linear NOTEARS formulation (Zheng et al., 2018).

NOTEARS formulates DAG structure learning as continuous optimization while enforcing an exact differentiable acyclicity constraint, thereby avoiding combinatorial graph search (Zheng et al., 2018). In DAG-HEART, the NOTEARS search space was further restricted using a predefined block-level directional mask. The model permitted miRNA-to-mRNA and DNA-methylation-to-mRNA dependencies, as well as dependencies among latent components within the miRNA and DNA-methylation blocks. Reverse mRNA-to-miRNA and mRNA-to-methylation connections and direct miRNA–methylation cross-block connections were excluded. The L_1_ regularization parameter for NOTEARS was 0.1, and graph optimization was performed for up to 30 iterations. Because the DAG was estimated from observational omics data, its edges were interpreted as directed statistical dependencies among latent components rather than as gene-level causal relationships. NOTEARS provides the continuous acyclicity framework on which this component of DAG-HEART is based.

For each permitted block relationship, the source-block latent representation was projected through the corresponding submatrix of the learned DAG and reconstructed into the feature space of the target omics block using the fitted PCA transformation. Four representations were subsequently retained for kernel construction as miRNA-to-mRNA, DNA-methylation-to-mRNA, miRNA-to-miRNA, and DNA-methylation-to-DNA-methylation. The same transformations learned from the training data were applied to validation and test samples.

### DAG-guided kernel integration

The magnitude of incoming and outgoing NOTEARS edge weights was used to quantify the relative contribution of each latent component. These latent importance values were propagated back to the molecular-feature space through the squared PCA loadings and were used to weight feature contributions during patient-similarity calculation.

For each of the four DAG-derived representations, weighted radial basis function (RBF) kernels were constructed to represent nonlinear patient-to-patient similarity. Kernel bandwidth was determined from the median nonzero squared Euclidean distance among training samples, and three bandwidth scales corresponding to 0.5, 1.0, and 2.0 times the base kernel parameter were considered. This procedure generated 12 candidate kernels within each fold. Kernel representations have previously been used to integrate heterogeneous genomic measurements by expressing each data source through sample-similarity relationships (Lanckriet et al., 2004).

The candidate kernels were integrated using centered kernel–target alignment. Kernel weights were estimated from the EA source-domain training samples by measuring alignment between each centered kernel matrix and the source-domain outcome similarity matrix. Negative alignment scores were truncated at zero, the remaining scores were normalized, and a smoothing term was applied to prevent individual kernels from dominating the integrated representation. The smoothing coefficient was set to 0.5. The resulting weighted kernel sum constituted the integrated patient-similarity matrix used for downstream TL.

Kernel principal component analysis (KPCA) was subsequently applied to the integrated kernel to obtain a compact nonlinear patient representation (Schölkopf et al., 1998). The minimum number of kernel components accounting for 90% of the nonnegative training-kernel eigenvalue sum was retained, with a maximum of 200 components. The fitted KPCA mapping was then applied to the source-domain samples, target-domain adaptation samples, validation samples, and held-out target-domain test samples. KPCA extends conventional principal-component analysis to nonlinear feature spaces through kernel eigenvalue decomposition (Schölkopf et al., 1998).

### Cross-validation and transfer-learning design

DAG-HEART was evaluated using repeated three-fold stratified cross-validation within the AA target cohort. Ten repetitions were performed using different random seeds. For each repetition, AA samples were divided into three folds while preserving the distribution of the fixed-time PFI classes. In each fold, one subset was held out for testing and the remaining AA samples were used for target-domain model development. From the target-domain training portion, up to five samples from each PFI class were randomly selected as labeled target samples for domain adaptation, and the remaining samples constituted the target-domain validation set. All available EA samples were used as the source-domain training cohort.

All representation-learning steps within each cross-validation fold, including block-wise scaling, PCA transformation, DAG estimation, kernel construction, kernel integration, and kernel PCA, were estimated before evaluation of the corresponding held-out AA test fold. Learning rates of 10^-3^, 3×10^-3^, and 10^-2^ and dropout rates of 0.1, 0.3, and 0.5 were evaluated as predefined model configurations. The overall experiment was therefore repeated across nine learning-rate/dropout combinations, with ten random-seed repetitions for each configuration.

TL was implemented using Classification and Contrastive Semantic Alignment (CCSA), a supervised domain-adaptation approach that learns a shared representation by jointly optimizing class discrimination and semantic alignment between labeled source- and target-domain samples (Motiian et al., 2017). CCSA was originally developed for supervised few-shot domain adaptation in settings where only a small number of labeled target-domain examples are available.

### Domain adaptation and data augmentation

The KPCA-derived representations were provided to a shared CCSA encoder containing one hidden layer with 100 units. The network jointly optimized a classification loss and contrastive semantic-alignment loss, with relative loss weights of 0.9 and 0.1, respectively. Models were trained using stochastic gradient descent with a momentum of 0.9 and a batch size of 20. Training was performed for up to 100 epochs with early stopping after 10 epochs without improvement on the target-domain validation data. For each fold, source–target semantic pairs were generated repeatedly to provide matched and mismatched outcome examples for contrastive domain alignment. This design follows the CCSA principle of preserving class discrimination while aligning semantically corresponding observations across domains (Motiian et al., 2017).

Two versions of the framework were evaluated. TL-only used the observed EA source and labeled AA target samples without target-domain augmentation. TL with DA additionally enabled synthetic balancing of the target domain during CCSA pair generation using the SMOTE-based augmentation procedure implemented in the model pipeline, with a maximum of five nearest neighbors. Synthetic minority over-sampling generates additional minority observations from local relationships among existing minority samples and was originally proposed to reduce class imbalance in supervised classification (Chawla et al., 2002).

For each fold, the classifier generated a probability for the positive fixed-time PFI class, which corresponded to progression-free status beyond the PFI time. A fold-specific classification threshold was selected exclusively from the target-domain validation set by maximizing the binary F1 score for this positive class. The selected threshold was then fixed and applied without further optimization to the corresponding held-out target-domain test samples. Predictions from the held-out folds were subsequently combined to obtain out-of-fold predictions for the AA target cohort for each random-seed repetition. Weighted F1 for the classifier-defined positive class was used solely as the validation-based criterion for selecting the fold-specific decision threshold, whereas support-weighted F1 was used to summarize held-out classification performance across both fixed PFI classes.

### Architectural and directional ablation analyses

To determine which components were responsible for the performance of DAG-HEART, four matched architectures were compared while retaining the same transfer-learning and target-domain augmentation procedure as a PCC-based model without DAG guidance, a DAG-guided PCC model, a nonlinear RBF–MKL–KPCA model without DAG guidance, and the complete DAG-HEART model combining DAG guidance with nonlinear kernel integration. The comparison therefore separated the contribution of directed representation learning from that of nonlinear kernel integration. AUROC, C-index, and weighted F1 were summarized over the same 10 repeated cross-validation runs. Module-specific effects were calculated as paired differences within the same PFI time setting and repeated run as DAG-guided minus non-DAG performance within PCC and kernel architectures, and kernel minus PCC performance with and without DAG guidance.

A second ablation evaluated the importance of the block-level directional prior. The proposed biologically motivated direction constraint was compared with a model in which the cross-omics directions were reversed and with a model in which no block-level directional prior was imposed. NOTEARS remained capable of estimating directed dependencies in the latter condition; only the predefined block-direction mask was removed. Matched differences between the proposed and comparator models were summarized within each PFI setting and repeated run. These repeated cross-validation contrasts were treated descriptively rather than as independent biological replicates, and no inferential P-values were assigned to differences among repeated runs.

### Stable molecular features and pathway analysis

Molecular interpretation focused on DAG-derived structural feature importance from the transfer-learning model with target-domain augmentation. For each fitted model, latent-node importance was defined from the combined absolute incoming and outgoing NOTEARS edge strengths and propagated to original molecular features through the squared PCA loadings. Structural importance was normalized to a mean of one within each fit and omics block. These values quantify a feature’s contribution to the learned DAG-guided representation and do not indicate the direction of its association with progression or establish a causal molecular effect.

For each PFI prediction, a feature’s recurrence was defined as the proportion of the 30 fitted models (10 repetitions × 3 folds) in which it ranked within the top 1% of structural importance within its omics block. A feature was operationally considered stable at a given PFI time when its top-1% recurrence was at least 50%. Cross-PFI setting importance and recurrence were then calculated by giving the 2-, 3-, 4-, and 5-year prediction tasks equal weight. Features stable in at least one PFI time setting were carried forward for pathway analysis.

Reactome over-representation analysis was conducted separately for mRNA, miRNA, and gene-level DNA methylation using clusterProfiler and Reactome gene sets obtained from the MSigDB C2:CP:REACTOME collection (Yu et al., 2012; Liberzon et al., 2011; Gillespie et al., 2022). clusterProfiler is an established framework for over-representation and functional enrichment analyses, and Reactome provides manually curated human pathway definitions. Pathways containing fewer than 10 or more than 500 genes were excluded. mRNA and gene-level methylation feature symbols were mapped to Entrez identifiers using org.Hs.eg.db, with the full set of successfully mapped measured genes serving as the corresponding omics-specific background. For miRNA, enrichment was performed on experimentally validated target genes from miRTarBase (Cui et al., 2025)

Over-representation testing was performed using a hypergeometric framework, and multiple-testing correction used the Benjamini–Hochberg procedure. For visualization, up to 10 pathways with nominal P<0.05 were displayed for each omics layer, while adjusted FDR values were retained to distinguish nominal enrichment from multiple-testing-corrected evidence. Cross-omics pathway summaries used the union of the top nominally enriched pathways, with FDR <0.05 indicated separately.

### Progression-risk stratification and PAM50 analyses

Clinical relevance was evaluated using unique-patient out-of-fold predictions from the AA target cohort. For each PFI setting, progression-risk estimates were averaged across the 10 repeated out-of-fold predictions for each patient. Progression risk was defined as one minus the mean event-free score. Patients were divided into high- and low-risk groups using the median progression risk calculated across the complete PFI time-specific AA cohort; this threshold was defined before any PAM50 stratification and was not recalculated within individual molecular subtypes.

Kaplan–Meier curves were generated for the PFI time-specific high- and low-risk groups. Continuous risk was additionally standardized within each PFI time setting and evaluated using a Cox proportional-hazards model, reporting the hazard ratio per one-standard-deviation increase in progression risk. Cox models used Efron’s method for handling tied event times, and the proportional-hazards assumption for the continuous risk term was assessed using Schoenfeld-residual-based tests.

PAM50 annotations were obtained using *TCGAbiolinks::PanCancerAtlas_subtypes()* (Colaprico et al., 2016), and intrinsic subtypes were interpreted according to established PAM50 breast-cancer classifications (Perou et al., 2000; Parker et al., 2009). Only TCGA sample type *01* (primary solid tumor) was considered. *Subtype_Selected* was preferentially used when available.

Progression-risk distributions were compared descriptively across Luminal A, Luminal B, HER2-enriched, and Basal-like tumors. To examine whether model-defined risk retained prognostic relevance within established molecular subtypes, restricted mean survival time (RMST) was calculated separately for each PAM50 subtype using the same PFI time-wide high/low-risk threshold. RMST was truncated at the corresponding model PFI time setting of 2, 3, 4, or 5 years, and the contrast was reported as RMST(low risk) minus RMST(high risk), such that positive values indicated longer average progression-free time in the low-risk group. RMST provides a summary of survival experience up to a prespecified PFI time setting without requiring proportional hazards for the group contrast (Royston & Parmar, 2013). Subtype-specific RMST contrasts were calculated only when both risk groups contained at least three patients.

## Data availability

The multi-omics and clinical outcome data used in this study were derived from The Cancer Genome Atlas breast invasive carcinoma cohort. TCGA molecular and clinical resources are available through the National Cancer Institute Genomic Data Commons (https://portal.gdc.cancer.gov/). MOTLAB is available at https://github.com/wan-lab/DAG-HEART.

## Acknowledgements

Research reported in this publication was supported by the U.S. National Science Foundation under Award Numbers 2500836 and 2614824, the National Cancer Institute of the National Institutes of Health under Award Number R03CA317707, the National Institute Of Alcohol Abuse And Alcoholism of the National Institutes of Health under Award Number R21AA032098, and the Office Of The Director, National Institutes Of Health of the National Institutes of Health under Award Number R03OD038391. This research was supported by the State of Nebraska through the Pediatric Cancer Research Group, part of the Child Health Research Institute. This work was also partially supported by the University of Nebraska Collaboration Initiative Grant from the Nebraska Research Initiative (NRI). The content is solely the responsibility of the authors and does not necessarily represent the official views of the funding organizations.

## Author contributions

M.B.: Data collecting and preprocessing, model development and optimization, data analysis and interpretation, manuscript preparation, editing and review. J.W.: Manuscript editing and review. S.W.: Study conceptualization, design, and supervision, manuscript editing and review.

